# Cross-protective immune responses induced by sequential influenza virus infection and by sequential vaccination with inactivated influenza vaccines

**DOI:** 10.1101/335281

**Authors:** Wei Dong, Yoshita Bhide, Federica Sicca, Tjarko Meijerhof, Kate Guilfoyle, Othmar G. Engelhardt, Louis Boon, Cornelis A.M. de Haan, George Carnell, Nigel Temperton, Jacqueline de Vries-Idema, David Kelvin, Anke Huckriede

**Author notes:** Address correspondence to Anke Huckriede. Present address: Kate Guilfoyle, Viroclinics Biosciences B.V., Rotterdam, The Netherlands.

## Abstract

Sequential infection with antigenically distinct influenza viruses induces cross-protective immune responses against heterologous virus strains in animal models. Here we investigated whether sequential immunization with antigenically distinct influenza vaccines can also provide cross-protection. To this end, we compared immune responses and protective potential against challenge with A(H1N1)pdm09 in mice infected sequentially with seasonal A(H1N1) virus followed by A(H3N2) virus or immunized sequentially with whole inactivated virus (WIV) or subunit (SU) vaccine derived from these viruses. Sequential infection provided solid cross-protection against A(H1N1)pdm09 infection while sequential vaccination with WIV, though not capable of preventing weight loss upon infection completely, protected the mice from reaching the humane endpoint. In contrast, sequential SU vaccination did not prevent rapid and extensive weight loss. Protection correlated with levels of cross-reactive but non-neutralizing antibodies of the IgG2a subclass, general increase of memory T cells and induction of influenza-specific CD4+ and CD8+ T cells. Adoptive serum transfer experiments revealed that despite lacking neutralizing activity, serum antibodies induced by sequential infection protected mice from weight loss and vigorous virus growth in the lungs upon A(H1N1)pdm09 virus challenge. Antibodies induced by WIV vaccination alleviated symptoms but could not control virus growth in the lung. Depletion of T cells prior to challenge revealed that CD8+ T cells, but not CD4+ T cells, contributed to cross-protection. These results imply that sequential immunization with WIV but not SU derived from antigenically distinct viruses could alleviate the severity of infection caused by a pandemic and may improve protection to unpredictable seasonal infection.

**Importance:** New influenza virus strains entering the human population may have large impact and therefore their emergence requires immediate action. Yet, since these strains are unpredictable, vaccines cannot be prepared in advance, at least not as long as there is no universal or broadly protective influenza vaccine available. It is therefore important to elucidate in how far immunization strategies based on currently available seasonal vaccines can provide at least some protection against newly emerging virus strains. Moreover, insight in the possible mechanisms of protection can guide the further development of pre-pandemic immunization strategies. Our study presents a vaccination strategy based on sequential administration of readily available seasonal whole inactivated virus vaccines which could be easily applied in case of a new pandemic. In addition, our study identifies immune mechanisms, in particular cross-reactive non-neutralizing antibodies and CD8+ T cells, which should be targeted by broadly protective influenza vaccines.

## Introduction

Influenza A virus (IAV) infections remain a worldwide public health threat. Influenza vaccination is the most reliable strategy to control annual epidemics and irregular pandemics [1]. Current inactivated influenza vaccines (IIV) primarily induce strain-specific antibodies against the two major virus surface proteins, hemagglutinin (HA) and neuraminidase (NA). However, these strain-specific antibodies cannot provide protection against antigenically drifted and antigenically shifted strains. When a pandemic strain emerges, it takes around six months to develop and distribute a new vaccine[2], which is too late for a vaccine to provide effective protection during the first pandemic wave. Thus, a cross-protective vaccine that could provide immediate protection against unpredicted influenza virus strains is urgently needed.

Live virus infection has been shown to provide some degree of cross-protection against A(H1N1)pdm09 infection in animal models[3][4][5][6][7][8] and in humans[9][10]. However, the exact mechanisms involved in cross-protection remain elusive. Cross-reactive antibodies against conserved regions of viral proteins, such as the HA stalk, the M2 ectodomain (M2e) and NP, induced by (sequential) live virus infection, correlate with cross-protection[3][11][12][13]. Some anti-HA stalk antibodies can directly neutralize influenza virus particles *in vitro*[14]. However, most of these antibodies target antigens that are expressed on the surface of infected cells and then provide cross-protection via a Fc receptor dependent mechanism[14][15][16].

Besides antibody responses, cross-reactive T cells induced by live virus infection have also been demonstrated to correlate with cross-protection[6][17][18]. Cytotoxic CD8 T cells can recognize internal, conserved epitopes across different virus strains. In animal models, CD8 T cells induced by live virus infection have been shown to prevent A(H5N1) or A(H1N1)pdm09 virus infection[19]. On the other hand, CD4 T cells specific for conserved epitopes have also been shown to provide protection against A(H1N1)pdm09 in mice[20][21]. These CD4 T cells could provide cross-protection through different mechanisms, including help for B cells, help for CD8 T cells and direct cytotoxic activity (reviewed in [22]). Furthermore, it has been demonstrated in humans that the presence of memory cross-reactive CD4 or CD8 T cells is correlated with cross-protection against A(H1N1)pdm09 or A(H7N9) virus infection[9][23][24].

Vaccination with trivalent inactivated influenza vaccine (IIV) did not provide protection against A(H1N1)pdm09 virus infection and was even found to be associated with enhanced disease in observational studies from Canada in humans[25][26][27][28][29]. In animal models, published studies indicate that vaccination with IIV could induce detectable levels of cross-reactive antibody against A(H1N1)pdm09 virus, yet, no cross-protection was observed[30][31][32]. The exception is a recent study showing that non-neutralizing antibody induced by IIV could cause activation of influenza-specific CD8 T cells by promoting antigen presentation[33]. If a broader immune response could be induced by the currently available influenza vaccines, it would benefit humans against novel virus infection.

Compared with a single virus infection, sequential infection with antigenically distinct live viruses was found to provide broader cross-protection[7][8][11]. This is because the second infection can cause a quick recall immune response to epitopes shared between the two viruses. It has been shown that sequential influenza virus infection can boost antibody responses to the shared HA stalk region[11][34].

Sequential immunization with antigenically distinct vaccines has also been used as a strategy to induce a broader immune response against influenza virus in animal models[35]. However, most of these studies were focused on the cross-protective immune response induced by genetically modified vaccines[36][37][38][39]. Little is known about the protective potential of sequential immunization with conventional inactivated vaccines derived from different seasonal influenza virus strains. In case of a pandemic, such a vaccination strategy could be a first means of intervention until a pandemic vaccine becomes available.

In this study, we assessed the cross-protective immune responses induced by sequential infection with A(H1N1) and A(H3N2) virus, or sequential immunization with whole inactivated virus (WIV) or subunit (SU) vaccine derived from these viruses in a mouse model. Sequential infection provided robust cross-protection which was mediated by non-neutralizing, cross-reactive antibody and CD8 effector memory T cells (TEM). Partial cross-protection was provided by sequential vaccination with WIV and was associated with CD8 central memory T cells (TCM), and to a minor extent, with cross-reactive antibodies. In contrast, sequential vaccination with SU vaccine induced low levels of cross-reactive serum antibodies and no T cell immunity against A(H1N1)pdm09, and did not provide cross-protection. These results imply that in case of a new pandemic, sequential immunization with WIV but not subunit vaccines derived from different seasonal virus strains could mitigate disease severity until a pandemic vaccine becomes available.

## Materials and Methods

### Virus and vaccines

Influenza virus strains A/Puerto Rico/8/34 (H1N1)(PR8), X-31, a reassortant virus derived from A/Aichi/68 (H3N2), and A/California/07/2009 (H1N1)pdm09 were grown in embryonated chicken eggs, and the virus preparations were titrated on MDCK cells and in mice. Whole inactivated virus vaccines was produced from PR8, X31 and X-181 (HA and NA proteins from A/California/7/2009 (H1N1)pdm09 and internal proteins from PR8) by treatment with β-propiolactone. PR8 subunit (SU) vaccine and X-31 SU were prepared from PR8 and X-31 WIV, respectively, as described before [40].

### Vaccination, challenge and sample collection

Female 6-8 weeks old CB6F1 mice) were purchased from Envigo, The Netherlands, and rested for at least one week. Mice were housed under SPF conditions in standard polycarbonate cages (5 animals per cage) with standard rodent bedding and cardboard cylinders as cage enrichment. Prior to the start of the experiment, animals were randomly allocated to the different treatment groups. All animal experiments were approved by the Central Committee for Animal Experiments CCD of the Netherlands (AVD105002016599). All experimental protocols were approved by the Animal Ethics Committee of the University Medical Center Groningen. Group sizes were determined using Piface software such that a power of at least 80% was reached.

Naive mice (n = 15) were immunized intramuscularly (i.m.) with 15 µg of PR8 WIV (containing around 5 µg of HA) or 5 µg of PR8 SU vaccine. Alternatively, mice were anesthetized and infected intranasally (i.n.) with a sublethal dose (10^3^ TCID_50_) of PR8 virus (live virus = LV). Four weeks after immunization or infection, mice were i.m. immunized with 15 µg of X-31 WIV or 5 µg of X-31 SU or i.n. infected with a sublethal dose of (10^3^ TCID_50_) X-31 virus. Mice injected twice with PBS i.m. with 28 days interval served as negative control (Table 1).

**Table 1.** Experimentaldesign for mouse experiment

| Groups | First immunization<br>(Day 0) | Second immunization<br>(Day 28) | Challenge<br>(D56) |
| --- | --- | --- | --- |
| 1 | PR8 WIV | X-31 WIV | H1N1pdm09 |
| 2 | PR8 SU | X-31 SU |  |
| 3 | PR8 LV* | X-31 LV |  |
| 4 | PBS | PBS |  |

\*LV = live virus

Four weeks after the second infection or immunization, 5 mice of each group were sacrificed for determination of infection- or vaccine-induced immune responses. The other 10 mice were anesthetized with isoflurane and challenged i.n. with 10^4.4^ TCID50 of A/California/7/2009 H1N1pdm09 in 40 µl PBS. Three days post infection, 5 mice were sacrificed for determination of immune responses and lung virus titers. The remaining 5 mice were monitored daily for body weight loss for two weeks. Body weight loss exceeding 20% was considered as humane endpoint.

On day 0 (before challenge) and day 3 post challenge, mice (n = 5 from each group) were sacrificed under isoflurane anesthesia. Serum, nose wash and bronchoalveolar lavage (BAL) were collected for further analysis. Lungs were perfused with 20 ml PBS containing 0.1% heparin through the heart right ventricle. Right lung lobes were collected, homogenized, snap-frozen and stored at −80°C for virus titration. The whole lung (day 0) or the left lung lobes (day 3) and the spleens were collected for lymphocyte isolation.

### Viral titer in lung

Lung tissue collected on day 3 post-challenge was weighed, homogenized in 1 ml of Episerf medium (Thermo Fisher Scientific) and then centrifuged at 1200 rpm for 10 minutes. Supernatants were collected, aliquoted, snap-frozen and stored at −80°C until use. Lung virus titers were determined by infection of MDCK cells in 96-well plates with serial dilutions of the lung supernatants as described before[40]. Viral titers, presented as log10 titer of 50% tissue culture infectious dose per gram lung (log_10_TCID_50_/g), were calculated based on the Reed-Muench method[41].

### Isolation of lymphocytes from lung and spleen

Spleens were homogenized in complete IMDM (with 10% FBS, 1% Penicillin-Streptomycin and 0.1% β-mercaptoethanol) using a GentleMACS dissociator (Miltenyi Biotec B, Leiden, The Netherlands). Cell suspensions were then forced through a cell strainer (BD Bioscience, Breda, The Netherlands) and treated with ACK lysis buffer (0.15 M NH_4_Cl, 10 mM KHCO_3_, 0.1 mM EDTA, pH 7.2) to remove erythrocytes.

PBS-perfused lungs for isolation of lymphocytes were homogenized using a GentleMACS dissociator (Miltenyi Biotec) and then digested by treatment with collagenase D (0.5 mg/lung) (Roche, Woerden, The Netherlands) in DMEM medium supplemented with 2% FBS at 37°C for 1.5 hour. The cell suspension was passed through a cell strainer. Lung lymphocytes in the filtered suspensions were enriched using lymphocyte density gradients (Sanbio, Uden, The Netherlands) according to the manufacturer’s protocol. The concentration of Granzyme B in lung homogenates was determined using Granzyme B Ready-SET Go ELISA kit (eBioscience) according to the manufacturer’s protocol.

### ELISA

For the detection of IgG, IgG1, IgG2a or IgA antibody against A(H1N1)pdm09 virus in serum and nasal wash, ELISA plates (Greiner, Alphen a/d Rijn, Netherlands) were coated with 0.3 µg/well of X-181 WIV, conserved M2e peptide (SLLTEVETPIRNEWGSRSNDSSD) or NP protein overnight at 37°C and ELISA assays were performed as described before[40]. For NA-specific ELISA, recombinant NA protein of A(H1N1)pdm09 was expressed and purified as described previously[42]. ELISA plates were coated with 0.1 µg/well of NA overnight at 4 °C and assays were performed as described[40].

### Pseudotype HA stalk neutralization assay

Pseudotyped viruses (PV) were produced by co-transfection of HEK293T/17 cells using the polyethylenimine transfection reagent (Sigma, cat: 408727). Lentiviral packaging plasmid p8.91 and vector pCSFLW bearing the luciferase reporter were transfected alongside the relevant HA glycoprotein genes in the plasmid pI.18 [43]. Parental PV were produced bearing the HA of A/California/7/09 (H1), or A/duck/Memphis/546/1974 (H11). A chimeric HA (cHA) consisting of the stalk from A/California/7/09 (H1) and head from A/duck/Memphis/546/1974 (H11) was also produced[44]. Pseudotype based microneutralisation assays (pMN) were performed as described previously [43]. Briefly, serial dilutions of serum were incubated with 1×10^6^ relative luminescence units (RLU) of HA bearing PV per well on a 96-well white plate for 1h at 37°C 5% CO_2_ in a humidified incubator. 1.5×10^4^ HEK293T/17 cells were then added per well and plates incubated at 37°C 5% CO_2_ for 48h before addition of Bright-Glo™ reagent (Promega) and measurement of luciferase activity. Analysis was performed using Graph-Pad Prism. Stalk-directed antibody presence was measured via antibody titers recorded against the cHA and both of its parental strains (H11 and H1 PV). No (or negligible) antibodies should be present against the exotic H11 HA, restricting neutralisation of the cHA PV to antibodies directed against the conserved H1 stalk of the cHA. Control antibodies used included mAb CR6261 (Crucell, Johnson and Johnson) and polyclonal antiserum Anti H11N9 (NIBSC).

### Intracellular cytokine staining

For IFNγ intracellular cytokine staining, lymphocytes (1.5-2 × 10^6^) from lung or spleen in complete IMDM medium were stimulated with CD28 (1 µg/ml, eBioscience), with or without X-181 WIV (10 µg/ml), overnight at 37°C in a 5% CO_2_ incubator. Protein transport inhibitor cocktail (eBioscience) was added for the last 4 hours of stimulation. Stimulated cells were stained with fluorochrome conjugated antibodies, including Alexa Fluor 700-antiCD3 (clone 17A2), FITC-antiCD4 (GK1.5), PerCP-cy5.5-antiCD8α (53-6.7), eFlour 450-antiCD62L (MEL-14), APC-antiCD44 (IM7) for 45 minutes. After surface staining, cells were stained with Fixable Viability Dye eFluor 780 (eBioscience) to identify dead cells. Cells were then fixed with IC fixation buffer (eBioscience) and permeabilized with permeabilization buffer (eBioscience) before intracellular staining with PE-cy7-antiIFNγ (clone XMG1.2) (all monoclonal antibodies from eBioscience). Samples were acquired on a BD LSRII and data were analyzed by Kaluza^®^ Flow Cytometry Analysis Software.

### ELISPOT and tetramer staining

Influenza NP-specific IFNγ-producing T cells were enumerated using a commercial mouse IFNγ ELISpot kit (MABTEC, The Netherlands) according to the manufacturer’s protocol. Briefly, splenocytes (2.5 x 10^5^/well) collected on day 0 post-infection were incubated with or without 5 µg/ml of the PR8 NP_366-374_ epitope (ASNENMDAM) in a pre-coated 96-well plate. After overnight incubation, IFNγ-producing T cells were detected using alkaline phosphatase-conjugated anti-mouse IFNγ antibody. Spots were developed with BCIP/NBT substrate and counted with an AID Elispot reader (Autoimmune Diagnostika GmbH, Strassberg, Germany). The number of antigen-specific IFNγ-producing cells was calculated by subtracting the number of spots detected in the unstimulated samples from the number in stimulated samples.

Tetramer staining for lung samples was performed as follows: isolated lung lymphocytes were incubated with A(H1N1)pdm09 NP_366-374_-tetramer-PE (containing the A(H1N1)pdm09 epitope ASNENME™) for 40 min and then stained with mouse anti-CD8α-PerCP-cy5.5 antibody for 40 min. Samples were acquired on a FACS Calibur^™^ BD II flow cytometer. Data were analyzed by Kaluza^®^ Flow Cytometry Analysis Software.

### Serum adoptive transfer

Mice were sequentially infected or sequentially immunized with WIV as described above and serum was collected on day 28 post second infection or immunization. Serum collected from mice that were immunized twice i.m. with A(H1N1)pdm09 WIV served as positive control. Pooled serum was tested by ELISA for presence of anti-A(H1N1)pdm09 antibodies. Naive mice (n = 5/group) received 200 µl of pooled serum by intraperitoneal injection one day before challenge with A(H1N1)pdm09 virus. On day 6 post challenge, lungs were collected for virus titration.

### CD4 and CD8 T cell depletion *in vivo*

For the T cell depletion study, mice were infected or vaccinated as described above and rested for 28 days. Groups of mice (n = 6/group) were injected with anti-CD4 T cell depletion antibody (200 µg/injection, GK1.5) or anti-CD8 T cell (200 µg/injection, YTS169). These antibodies were given i.p on day -1, 1 and 3 of A(H1N1)pdm09 virus (10^4.4^ TCID_50_) challenge. On day 6 post challenge, lungs were collected for virus titration. Spleens were collected to confirm the depletion of T cells.

### Statistics

Mann-Whitney U test was used to determine the differences between read-outs of two different groups. Statistical analyses were performed using GraphPad Prism version 6.01 for Windows. GraphPad Sofware, La Jolla, California, USA www.graphpad.com. P < 0.05, 0.01, 0.001 were considered as significantly different and were denoted by *, **, ***, individually.

## Results

### Sequential infection, WIV and SU vaccination show different levels of cross-protective capacity against H1N1pdm09 influenza virus infection

To investigate the cross-protective immune response induced by sequential infection or vaccination with antigens from different influenza virus strains, we sequentially infected mice with PR8 and X-31 influenza virus or sequentially vaccinated mice with WIV or SU vaccines derived from these viruses. These viral strains were selected to reflect a heterosubtypic exposure history in humans. The cross-protective capacity of sequential infection or sequential immunization was determined by challenging the mice with virus A/California/7/2009 (H1N1)pdm09.

After A(H1N1)pdm09 virus challenge, mice in the sequential SU vaccination group showed similar weight loss as mice in the PBS control group and developed severe symptoms, necessitating euthanasia on day 6 or 7 post challenge (Fig. 1 A, B). Mice that were sequentially vaccinated with WIV showed a similar trend of weight loss as mice in the PBS control group until day 6 post infection. Yet, from day 7 post infection onwards, WIV immunized mice recovered and none of the mice reached the humane endpoint. In the sequential infection group, mice showed no or only minor weight loss after challenge and none of them needed to be sacrificed.

**Figure 1.**
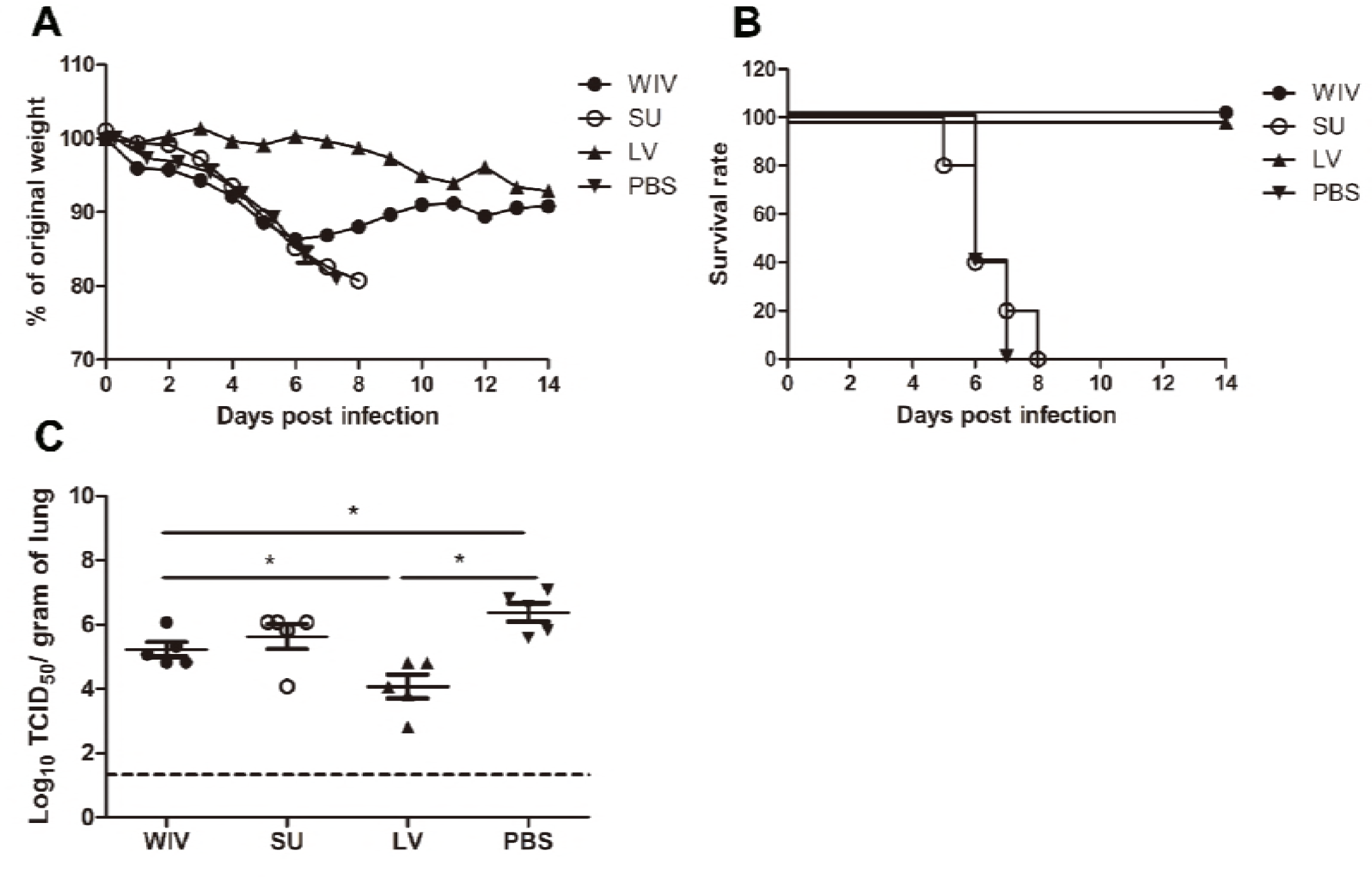
Weight loss and survival rate of immunized mice after A(H1N1)pdm09 virus challenge. Naïve mice (n=10) were sequentially infected with sublethal doses of two different strains (PR8 and then X31) of live virus (LV) with 28 days interval or were sequentially immunized with vaccines (WIV, SU) derived from these virus strains and then challenged with virus A/California/7/2009 (H1N1)pdm09. After challenge, mice (n=5) were monitored daily for weight loss (A) and survival (B) for a period of 14 days. On day 3 post-challenge, lung virus titers in 5 mice/group were determined by titration on MDCK cells (C). * p<0.05, Mann-Whitney U test. The dashed line represents the limit of detection.

On day 3 post-challenge, lung virus titers in the sequential SU vaccination group did not differ significantly from those in the PBS control group (Fig. 1C). In the sequential WIV vaccination group, lung virus titers were decreased by 0.9 log_10_ as compared to the PBS group (p = 0.03). Sequential infection resulted in a significant decrease of the lung virus titer by 2 log_10_ relative to the control group (p = 0.015).

These data demonstrate that sequential immunization with WIV, although being less effective than sequential infection with live virus, provided a certain level of cross-protection against heterologous infection. In contrast, sequential SU vaccination did not provide cross-protection.

### Sequential infection, WIV and SU vaccination induce distinct cross-reactive antibody immune responses

To explore the immune mechanisms involved in protection from weight loss and lung virus growth upon challenge, A(H1N1)pdm09 cross-reactive antibody responses induced by sequential infection with PR8 and X-31 or immunization with PR8 and X-31 derived vaccines were determined.

Sequential infection induced around 20 times more cross-reactive IgG antibody than sequential WIV vaccination and approximately 75 times more cross-reactive IgG antibody than sequential SU vaccination (p < 0.0001) (Fig. 2A). With respect to the subtype profile of the IgG antibodies, sequential infection and WIV vaccination induced a Th1-type antibody response.

**Figure 2.**
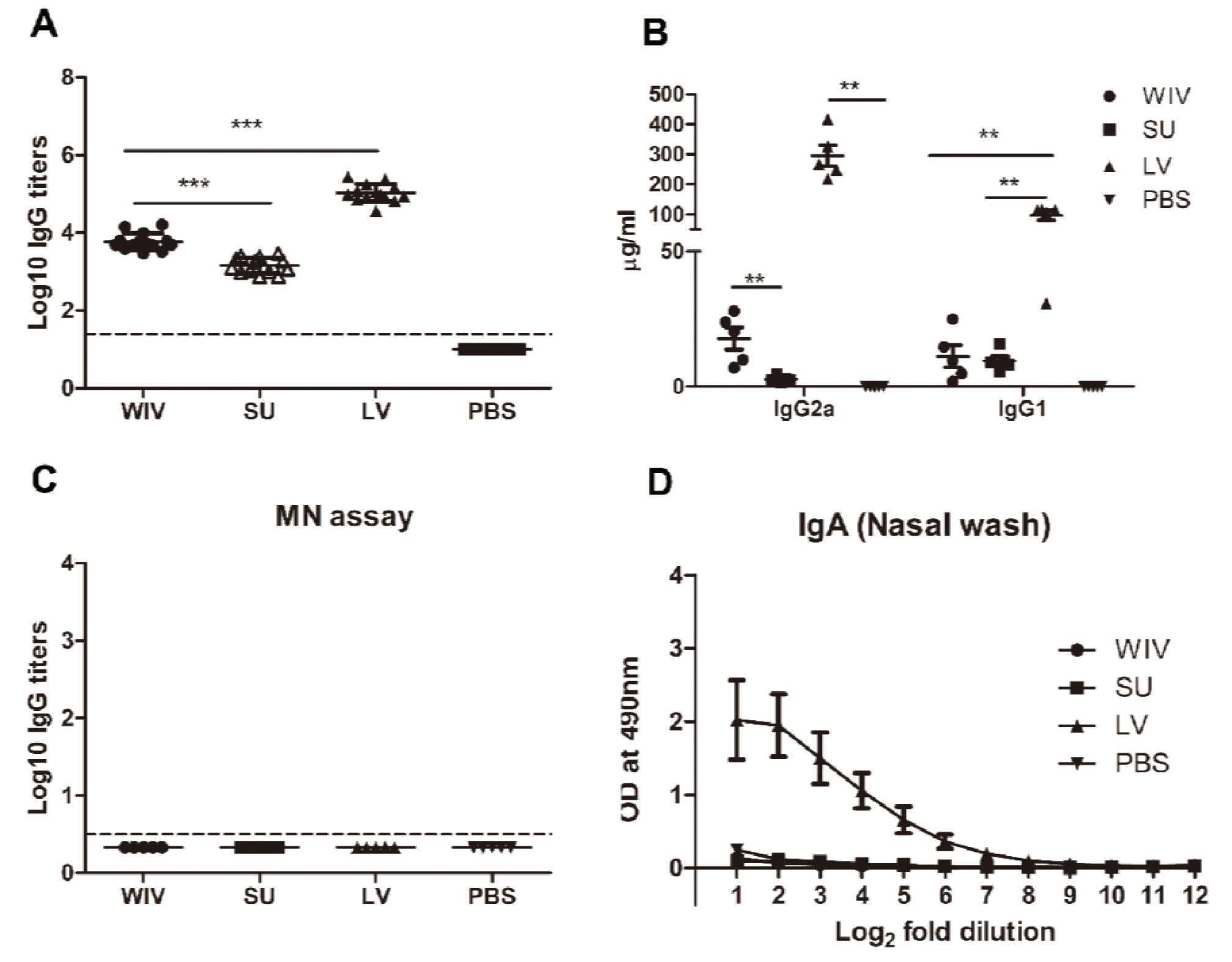
Cross-reactive antibody immune response induced by sequential infection or immunization. On day 28 post the second infection or immunization, serum samples and nasal washes were collected from the mice described in the legend to Fig. 1. Anti-H1N1pdm09-specific IgG (A; n=15), IgG2a and IgG1 (B; n=5) antibodies in serum samples were detected by ELISA. Microneutralization assay was used to determine the neutralizing ability of these antibodies towards A(H1N1)pdm09 virus (C; n=5). Anti-H1N1pdm09 IgA antibody levels in nasal washes were determined by ELISA (D; n=5). Data of individual animals (A, B, C) are depicted or mean values ± SEM (D) are given, **, p<0.01, ***, p<0.001. Mann-Whitney U test. The dashed line represents the limit of detection.

The average ratio of serum IgG2a to IgG1 concentration was 3 for mice sequentially infected by live virus, compared with 1.5 induced by sequential WIV vaccination. In contrast, sequential SU vaccination induced a similar amount of IgG1 antibody as induced by sequential WIV vaccination but no IgG2a (Fig. 2B). However, cross-reactive antibodies, irrespective of whether induced by sequential infection or immunization, did not neutralize A(H1N1)pdm09 virus (Fig. 2C). With respect to mucosal antibodies, only sequential infection was found to induce cross-reactive IgA antibody against A(H1N1)pdm09 virus in the nose (Fig. 2D).

In order to reveal the target protein(s) of the observed cross-reactive antibodies we first performed a pseudovirus-based assay to detect antibodies to the HA stalk domain. This assay uses a chimeric HA as antigen, with an H11 globular head, and an H1 stalk. The chimeric HA pseudovirus particles were effectively neutralized by the CR6261 mAb control which binds to the H1 stalk. However, no antibodies reacting with the H1 stalk were observed in any of the experimental groups (data not shown). Next, we examined anti-NA antibodies against A(H1N1)pdm09 virus. The mice from the sequential infection group and 4 out of 5 mice from the WIV vaccination group developed anti-NA antibodies, while only 2 out of 5 mice from the sequential SU vaccination group did so and levels of anti-NA antibody were low (Fig. 3A). Next, anti-M2e antibody titers were determined by coating conserved M2e peptide onto 96-well ELISA plate. Anti-M2e antibodies were only found in the sequential infection group (Fig. 3B). We also analyzed the presence of cross-reactive antibodies against conserved internal proteins in serum using recombinant NP from HK68 (H3N2), which shows 90% of sequence homology with NP from A(H1N1)pdm09. Sequential infection and WIV vaccination induced similar though somewhat variable amounts of anti-NP antibodies (Fig. 3C). As expected, no anti-NP antibody was found in the sequential SU vaccination group.

**Figure 3.**
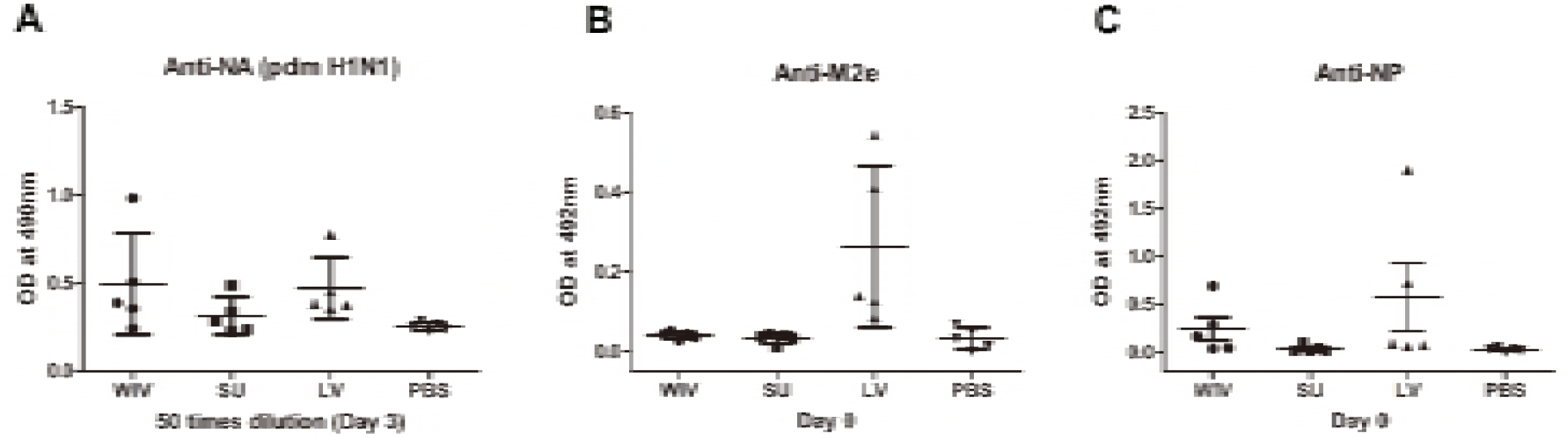
Cross-reactive antibodies against conserved proteins induced by sequential infection or immunization. Mice were primed and boosted as described in the legend to Fig 1. Serum samples were collected 28 days post boost (day 0) or 3 days post-challenge (day 3). (A) Antibodies against A(H1N1)pdm09 NA protein on day 3 post-challenge were determined by ELISA. Anti-M2e (B) and Anti-NP (C) antibodies titers were determined by ELISA. Data represent mean values ± SEM.

These data indicate that sequential infection induced broader and higher amounts of cross-reactive non-neutralizing antibodies than sequential WIV vaccination, while SU vaccination induced only antibodies against hemagglutinin and to a limited extent against neuraminidase. Moreover, responses to live virus and WIV were dominated by IgG2a while responses to SU consisted exclusively of IgG1 antibodies.

### Sequential infection, WIV and SU vaccination induce different memory T cell immune response

Apart from cross-reactive antibody response, cellular immune responses also play an important role in cross-protection. We first evaluated the overall memory T cell responses in spleen and lungs from mice after sequential infections or vaccinations. None of these immunization strategies could significantly enhance the number of memory CD4+CD44+ T cells (p = 0.28) (Fig. 4A). However, numbers of memory CD8+CD44+ T cells were significantly enhanced in spleen (p = 0.028) and lung (p = 0.015) of sequentially infected mice compared with mice of the unvaccinated group (Fig. 4B). Also, sequential WIV vaccination enhanced memory CD8+CD44+ T cell numbers, however, only in spleen was significance reached (p = 0.02) (Fig. 4B). No increase in the number of memory CD8 T cells was found in the sequential SU vaccination group. Interestingly, while the CD8 memory T cell population in sequentially infected mice consisted of CD62L negative TEM as well as CD62L positive TCM, the majority of memory CD8 T cells from the sequential WIV vaccination group was CD62L positive (Fig. 4C). These data indicate that sequential infection and sequential immunization with WIV are capable of stimulating CD8 memory responses while immunization with SU is not.

**Figure 4.**
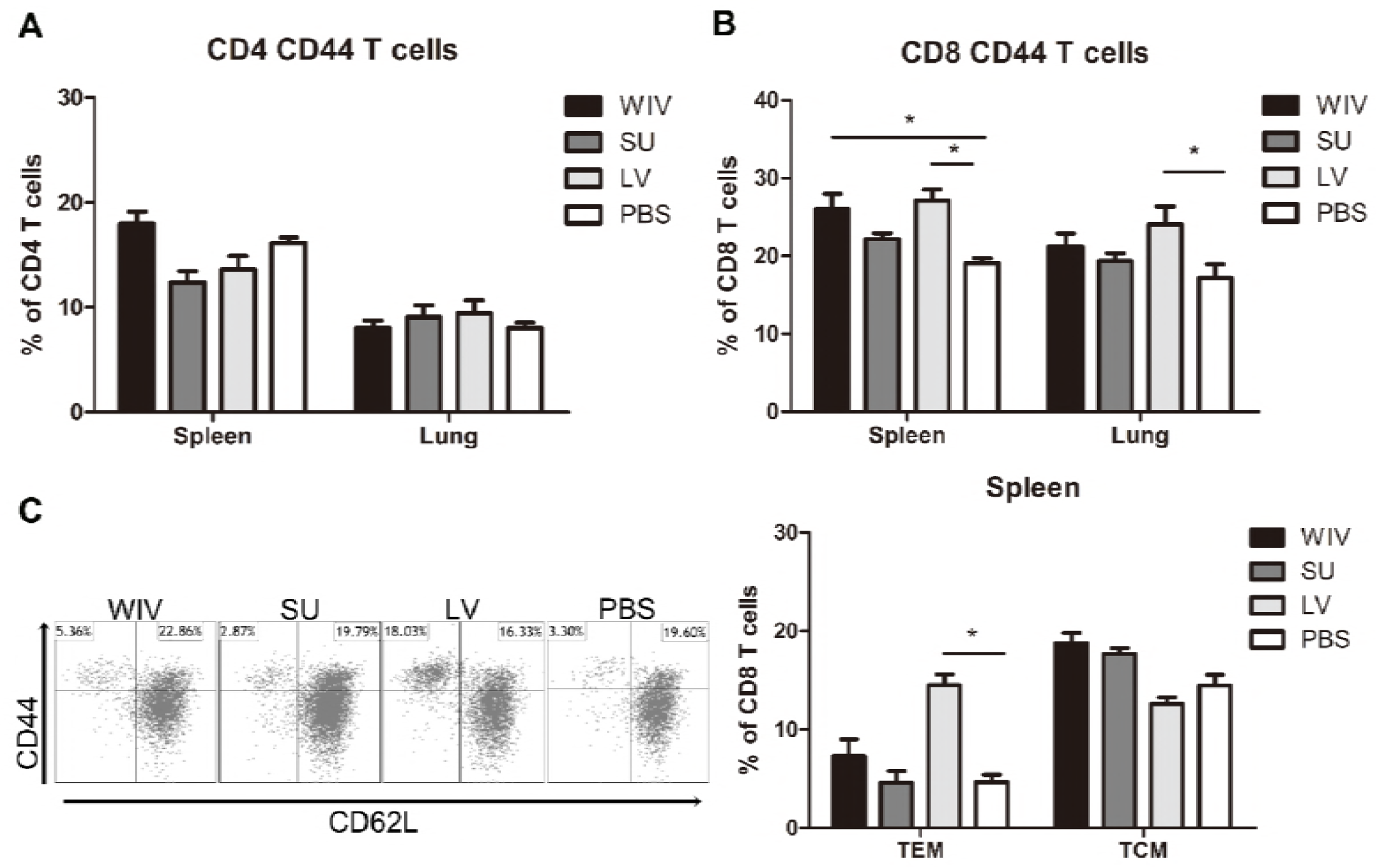
Memory T cell immune responses after sequential infection or immunization. Of the mice described in the legend to Fig. 1, 5 animals/group were sacrificed 28 days after the second infection or immunization and spleen and lung were collected. (A) CD4+CD44+ and (B) CD8+CD44+ memory T cells in spleen and lung were determined by flow cytometry. (C) CD8+CD44+CD62L-effector memory T cells (TEM) and CD8+CD44+CD62L+ central memory T cells (TCM) in spleen. Left: representative dot plots depicting CD44 and CD62L expression on spleen CD8 T cells. Right: percentages of spleen CD8 TEM and TCM + SEM. (n=4 or 5 per group, representative of two experiments, Mann-Whitney U test, *, p<0.05).

For detection of influenza specific T cells, splenocytes from sequentially infected or sequentially immunized mice were stimulated overnight with WIV, and IFNγ production was assessed by intracellular cytokine staining. In live virus infected mice, percentages of IFNγ producing CD4+ and CD8+ memory T cells in spleen and lung were significantly higher than in mock immunized mice (Fig. 5A, p < 0.05). Moreover, around 90% of these IFNγ-producing CD8 T cells were effector memory cells (data not shown). Also in WIV immunized mice, enhanced percentages of IFNγ positive CD4+ and CD8+ memory T cells were found, yet lower than in the LV group. Significance as compared to PBS control animals was reached only for CD4+ T cells in spleen.

**Figure 5.**
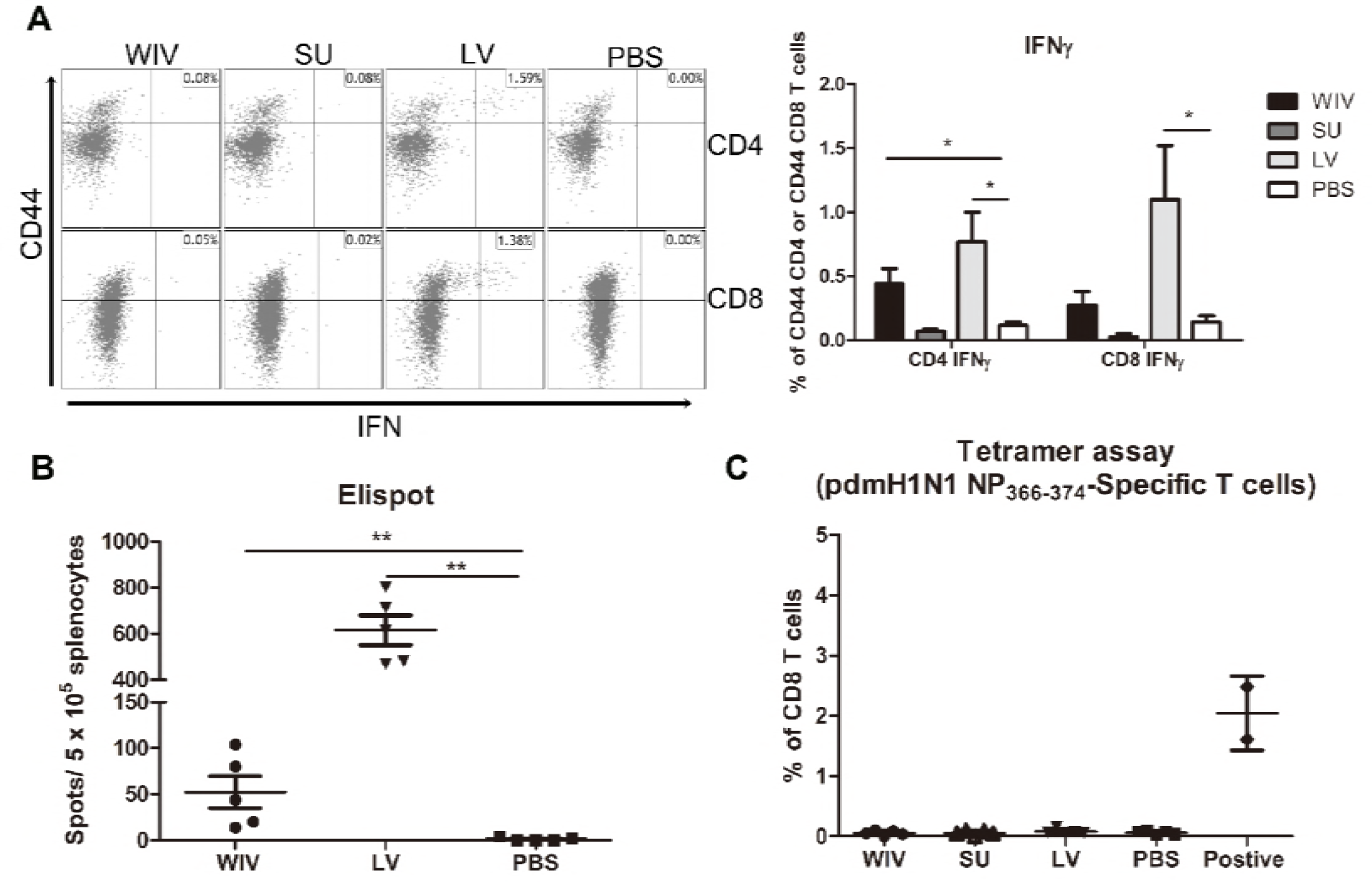
Influenza-specific T cell immune responses induced by sequential infection or immunization. (A) Splenocytes harvested on day 28 post the second infection/vaccination, were stimulated with A(H1N1)pdm09 WIV and anti-CD28 overnight in presence of protein transport inhibitor. Presence of intracellular IFNγ in CD4+CD44+ and CD8+CD44+ T cells was analyzed by flow cytometry. Left: representative dot plots of stimulated CD4 or CD8 T cells stained for CD44 and IFNγ. Right: percentages of IFNγ-producing cells among CD4+CD44+ and CD8+CD44+ T cells. (n=4 or 5, representative of two experiments, Mann-Whitney U test, *, p<0.05). (B) On day 28 post the second infection/immunization, NP366-374 of PR8 virus was used to stimulate mouse splenocytes and IFNγ-producing CD8 T cells were enumerated by ELISPOT. (n=5, Mann-Whitney U test, **, p<0.01). (C) A(H1N1)pdm09 NP366-374-specific CD8 T cells in spleens of infected/immunized mice (n=5) were determined by tetramer assay. Lymphocytes from the blood sample of mice (n=2) infected with A(H1N1)pdm09 virus served as positive control.

The influenza-specific CD8 T cells induced by infection or immunization were also enumerated by ELISPOT after stimulation of splenocytes with NP_366-374_ peptide (ASNENMDAM) from PR8 virus (the epitope presents in PR8 as well as X-31 virus). NP-specific CD8 T cells were detected in the WIV and the sequential infection group, but numbers were around 12 times higher in the latter (Fig. 5B, p = 0.008). Next, we assessed the cross-reactivity of these NP-specific CD8 T cells to A(H1N1)pdm09 NP by staining with tetramers containing the ASENENME™ epitope (from A(H1N1)pdm09 virus). No tetramer positive CD8 T cells were observed in these groups of mice (Fig. 5C) while tetramer positive cells were readily detected in blood of mice infected with A(H1N1)pdm09 virus.

### Serum antibodies induced by sequential infection are sufficient to provide cross-protection but antibodies induced by WIV vaccination are not

Our data show that sequential infection and sequential immunization with WIV could provide protection against severe symptoms upon infection with an A(H1N1)pdm09 virus. To determine the contribution of cross-reactive antibodies against A(H1N1)pdm09 virus challenge, serum from sequentially virus infected, WIV vaccinated or PBS control mice was passively transferred to naive mice one day before A(H1N1)pdm09 virus challenge. Serum from mice vaccinated with WIV derived from A(H1N1)pdm09 virus served as positive control.

Mice receiving serum from mice immunized with A(H1N1)pdm09 WIV (positive control, neutralizing titer 330) via adoptive transfer did not show weight loss upon A(H1N1)pdm09 virus challenge (Fig. 6A) and lung virus titers in these animals were decreased by more than 2 logs compared to the titers in the PBS control group (Fig. 6B, p < 0.01). Similarly, mice receiving serum from the sequential infection group showed no or only mild weight loss. Interestingly, despite the fact that the transferred serum did not contain any neutralizing antibodies, lung virus titers in this group were decreased to the same low level as in mice which had received serum from A(H1N1)pdm09-immunized mice containing neutralizing antibodies. Also serum from the sequential WIV vaccination group provided partial protection; 4 out of 5 mice receiving this serum showed no or mild weight loss, while one mouse went down quickly. Yet, lung virus titers in the WIV vaccination group, though slightly lower, did not differ significantly from those in PBS-treated controls (p = 0.22) (Fig. 6B).

**Figure 6.**
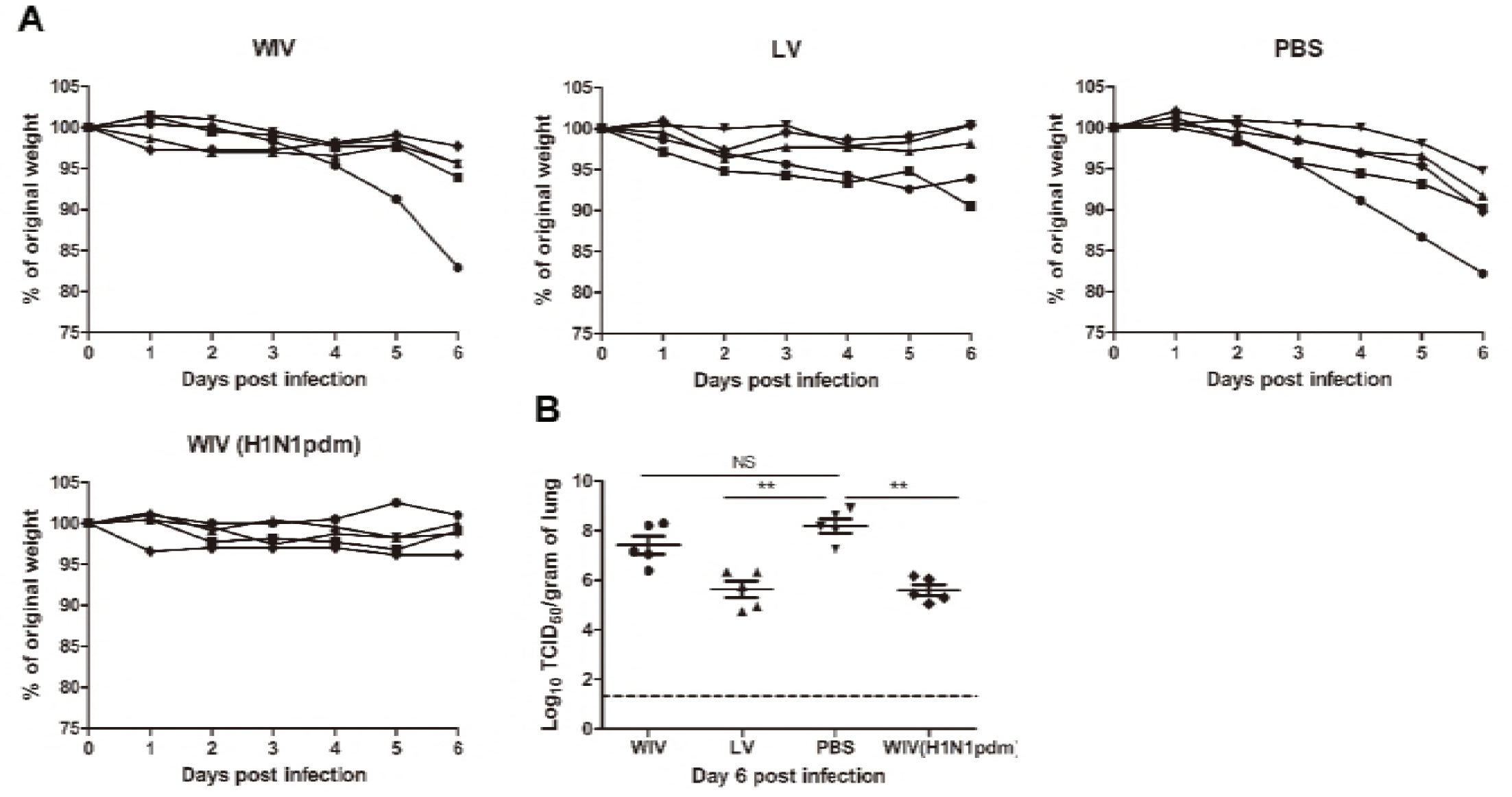
The cross-protective potential of antibodies induced by sequential infection or immunization. Mice (n=5) were primed with PR8 virus (10^3^TCID_50_) or PR8 WIV (15µg) and boosted with X-31 virus (10^3^TCID_50_) or X-31 WIV (15µg). Mice primed and boosted with PBS served as negative control and mice primed and boosted with A(H1N1)pdm09 WIV (15µg) served as positive control. Sera from these mice were collected 4 weeks after boost, pooled and injected intonaïve mice one day before challenge with A/California/7/2009 (H1N1)pdm09 virus. Body weight loss (A) was monitored daily for 6 days. Virus titers in the lung tissue (B) on day 6 post-challenge were determined by titration on MDCK cells. **, p<0.01, Mann-Whitney U test. The dashed line represents limit of detection. NS, not significant.

These data indicate that non-neutralizing antibodies induced by sequential infection were as effective as neutralizing antibodies induced by A(H1N1)pdm09 WIV vaccination in providing protection against A(H1N1)pdm09 virus challenge. However, non-neutralizing antibody induced by sequential WIV vaccination were not sufficient to provide full cross-protection.

### Memory T cells induced by sequential live virus infection or WIV vaccination are involved in cross-protection against A(H1N1)pdm09 virus challenge

To determine the contribution of T cell immune responses to cross-protection against A(H1N1)pdm09 virus infection, we used CD4 or CD8 specific antibodies to deplete T cells before and during A(H1N1)pdm09 challenge. On day 6 post-challenge, we confirmed that 95% of CD8 T cells or 96% of CD4 T cells in mice spleen were depleted by this treatment (data not shown).

Mice in the PBS mock vaccination group, no matter whether treated with PBS, CD4 depletion antibody or CD8 depletion antibody, showed continuous weight loss after A(H1N1)pdm09 challenge (Fig.6A, PBS) and displayed the same virus titers in lung tissue on day 6 post-infection (Fig. 6B, PBS). In contrast, mice in the sequential infection group were protected from weight loss and showed low or undetectable lung virus titers (Fig. 7A, LV). Depletion of CD4 T cells in these mice had no effect on protection. Depletion of CD8 T cells in the sequential infection group had some effect on protection from weight loss; on day 6 post A(H1N1)pdm09 virus challenge 3 out of 6 mice had lost > 6.5 % weight while in non-depleted mice the most severe weight loss was 2.1% and was observed in a single mouse only (Fig. 7A, LV). In addition, lung virus titers were about 1.5 log_10_ higher in the CD8-depleted mice than in non-depleted control mice of the sequential infection group; yet, virus titers were still significantly lower than in non-immunized mice. In the WIV vaccination group, depletion of CD4 or CD8 T cell did not significantly alter the weight loss compared with mock depletion but a strong trend towards less weight loss was observed in mice depleted for CD4 T cells as compared to non-depleted mice of this group (P = 0.054, Fig. 7A, WIV). Depletion of CD4 T cells decreased and depletion of CD8 T cells increased lung virus titers by about 1 log as compared to non-depleted animals on day 6 post challenge but these trends did not reach statistical significance (Fig. 7B, WIV). Moreover, virus titers in WIV-immunized CD8 T cell-depleted mice were of the same magnitude as those in the PBS mock vaccination group.

**Figure 7.**
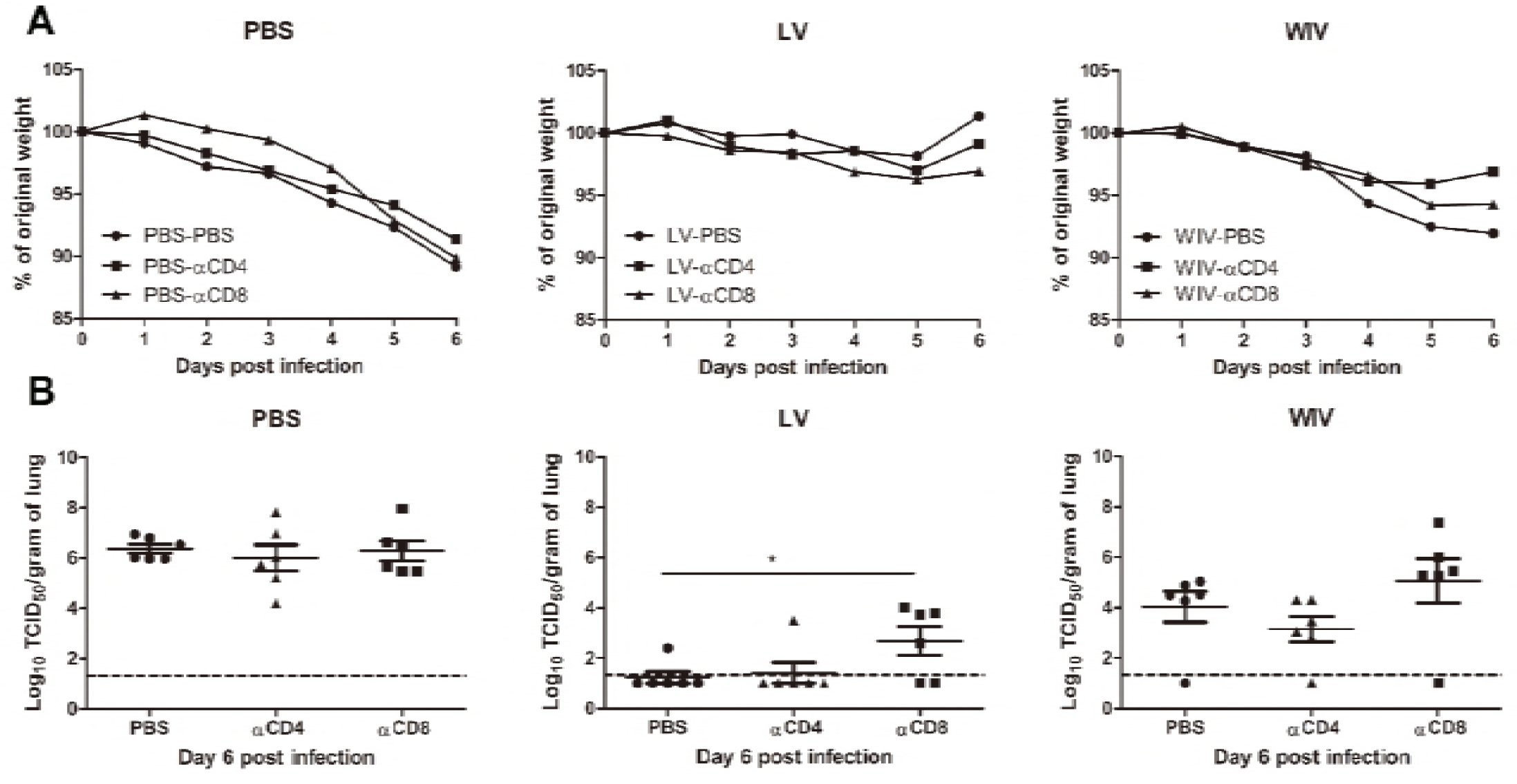
The cross-protective potential of CD4 T cells and CD8 T cells induced by sequential infection or immunization. Mice were primed with PR8 virus (10^3^TCID_50_) or PR8 WIV (15µg) and then boosted with X-31 virus (10^3^TCID_50_) or X-31 WIV (15µg). Mice primed and boosted with PBS served as control. Anti-CD4, anti-CD8 T cell depletion antibody or PBS were injected intraperitoneally intomice on day -1, 1 and 3 of A(H1N1)pdm09challenge. Weight loss (A) was monitored for 6 days and lung virus titers (B) were determined on day 6 post-infection by titration on MDCK cells. *, p<0.05, Mann-Whitney U test. The dashed line represents limit of detection.

These data above suggests that CD4 memory T cells were most likely not involved in cross-protection while CD8 memory T cells induced by sequential infection or WIV immunization contributed decisively to cross-protection.

## Discussion

To determine whether sequential immunization with antigenically distinct traditional vaccines could provide cross-protection, mice were sequentially immunized with WIV or SU vaccines derived from PR8 and X-31 viruses and then challenged with an A(H1N1)pdm09 virus. Another group of mice was sequentially infected with sublethal doses of PR8 followed by X-31 prior to A(H1N1)pdm09 virus challenge. We demonstrate that sequential infection provided solid cross-protection which was correlated with cross-protective antibodies and CD8 TEM cells. Sequential vaccination with WIV provided partial cross-protection which also correlated with induction of cross-reactive antibodies and CD8 T cells. Yet, sequential SU vaccination did not provide cross-protection.

Neither sequential infection nor sequential immunization resulted in induction of antibodies capable of neutralizing A(H1N1)pdm09 virus. Yet, substantial amounts of cross-reactive non-neutralizing antibodies were induced. Previous publications have shown that non-neutralizing antibodies, for example anti-HA stem antibodies, can be induced by sequential infection with antigenically distinct viruses and may provide cross-protection against A(H1N1)pdm09 influenza virus infection[11][12]. In contrast to these findings, no anti-HA stem antibodies were found in this study. This may be due to the fact that the two virus strains (PR8 and X-31) used for infection/immunization belong to two different phylogenetic groups. The HA-stem regions from PR8 and X-31 virus show low similarity, which might have impaired boosting of HA-stem reactive B cells induced by PR8 through exposure to X-31. Nevertheless, we found cross-reactive antibodies against other conserved proteins in this study. Anti-M2e, anti-NP and anti-NA antibodies were induced by sequential infection and, although to a lesser extent, by sequential WIV immunization. In contrast, sequential SU immunization induced only very moderate amounts of anti-NA antibodies cross-reactive with A(H1N1)pdm09 virus.

Since no neutralizing antibodies were found, the cross-reactive but non-neutralizing antibodies likely are the reason for the cross-protection observed in the serum adoptive transfer experiment. Non-neutralizing antibodies can provide cross-protection via Fc receptor dependent mechanisms (reviewed in [45]). Interestingly, control of lung virus growth by non-neutralizing antibodies evoked by sequential infection with PR8 and X-31 was as effective as by neutralizing antibodies evoked by A(H1N1)pdm09 WIV. Even in absence of antigen-specific T cells, neutralizing antibodies are thus not crucial for protection, suggesting that non-neutralizing antibodies maybe more important for cross-protection than generally thought. Interestingly, recent studies revealed that in humans antibodies cross-reacting with different influenza virus strains are common and that these antibodies are effectively enhanced by vaccination with seasonal influenza vaccines[46][47].

Hillaire et al and Guo et al have shown that one dose of serum from virus-infected animals could not provide cross-protection against A(H1N1)pdm09 virus infection in mice[6][17], while Fang et al have shown that four doses of serum could provide cross-protection[3]. These studies imply that the amount of non-neutralizing cross-reactive antibodies may also play an important role in cross-protection. In the present study, cross-reactive antibody titers evoked by sequential WIV immunization were 20-fold lower than those evoked by sequential infection. We thus speculate that antibodies induced by WIV immunization, though in principle cross-protective as indicated by our data, were not present in sufficient amounts to confer complete protection.

Although sequential infection and sequential WIV immunization induced virus-specific IFNγ-producing CD4 T cells, depletion of CD4 T cells in this study did not influence the cross-protection, neither in the sequential infection group nor in the sequential WIV vaccination group. These results contrast with previous findings which indicate that CD4 T cells might play a role in cross-protection[22][6][17]. Hillaire et al reported that naïve mice that received T cells (a mixture of CD4 and CD8 T cells) induced by a single A(H3N2) (HK68) virus infection acquired better cross-protection against A(H1N1)pdm09 virus infection than naïve mice that received purified CD8 T cells only[6]. Another study by Guo et al reported that depletion of CD4 T cells induced by a single X-31 virus infection impaired the cross-protection against A(H1N1)pdm09 virus infection in mice[17]. In this study, not only CD4 T cells, but also robust cross-reactive antibodies and CD8 T cell immune responses were induced by sequential infection. These antibodies or CD8 T cells alone could significantly reduce the virus titer in mice lung in the absence of CD4 T cells. We conclude that CD4 T cell are not essential for cross-protection against A(H1N1)pdm09 during infection in this mouse model.

CD8 T cells play an important role in cross-protection. In the present study, depletion of CD8 T cells induced by sequential WIV immunization resulted in lung virus titers similar to those in PBS mock vaccinated mice, implying that CD8 T cells are important for cross-protection induced by sequential WIV immunization. These results agree with those reported by Furuya et al who showed that WIV (prepared by γ-irradiation) did not provide cross-protection against heterologous virus infection in mice defective in CD8 T cells [48]. Another study by Budimir et al also has shown that depletion of CD8 T cells induced by 2 doses of WIV abolished the cross-protection against heterologous virus challenge[49]. Depletion of CD8 T cells in the sequential infection group prior to A(H1N1)pdm09 challenge had a significant though moderate effect on lung virus titers. This result implies that in the sequential infection group CD8 T cells do play a role in cross-protection, but team up with other mechanisms, eg antibodies (Fig. 5), to provide full protection. Our findings are also in line with previous publications which demonstrate that CD4 T cells or antibody immune responses are required to cooperate with CD8 T cells for providing optimal cross-protection in live virus infected mice[18][17][50].

The tetramer experiment indicates that PR8 NP_366-374_ epitope-specific CD8 T cells elicited by PR8 and boosted by X-31 virus or WIV could not recognize the corresponding A(H1N1)pdm09 NP_366-374_ epitope. This result is in line with previous findings demonstrating that X-31 NP_366-374_ epitope cannot be recognized by A(H1N1)pdm09 NP-specific CD8 T cells[51]. However, Guo et al have reported that influenza NP and PA proteins from PR8 and A(H1N1)pdm09 virus share many conserved epitopes[51]. It is possible that influenza-specific CD8 T cells against these shared conserved epitopes induced by sequential infection or WIV immunization provided cross-protection against A(H1N1)pdm09 influenza virus infection.

Different phenotypes of memory CD8 T cells show different capacities in cross-protection, for example Wu et al have shown that CD8 TCM induced by influenza virus infection are not required for cross-protection[18]. In the present study, we found that sequential infection mainly induced CD8 TEM. This result is in line with previous findings in mice and humans reporting that a single influenza infection predominantly induces influenza-specific CD8 TEM cells[52][53]. CD8 TEM have been shown to be associated with a fast recall immune response to the infection site, thus providing immediate cross-protection[52]. Interestingly, we found that sequential WIV immunization was more likely to induce CD8 TCM. These cells have shown high proliferation ability in secondary lymphoid organs but to provide delayed cross-protection[54]. Thus, we propose that CD8 TEM in lung and spleen induced by sequential infection provided immediate local antiviral effects, resulting in solid cross-protection. In contrast, CD8 TCM in spleen induced by sequential WIV immunization provided delayed antiviral effects in the lung, resulting in partial cross-protection.

In summary, sequential infection with antigenically distinct viruses provided solid cross-protection against A(H1N1)pdm09 virus infection. Yet, sequential immunization with antigenically distinct SU failed to provide cross-protection. Intriguingly, sequential immunization with antigenically distinct WIV provided partial cross-protection by a mechanism involving cross-reactive but non-neutralizing antibodies as well as CD8+ T cells. These results imply that sequential immunization with WIV prepared from antigenically distinct viruses could be used to alleviate the severity of virus infection if a new pandemic occurs.

## Acknowledgements

The authors would like to thank Meiling Dai, Utrecht University, Utrecht, The Netherlands, for technical assistance and David A. Price, Institute of Infection and Immunity, Cardiff University School of Medicine, Cardiff, United Kingdom, for a gift of tetramers. This research was funded by the European Union Seventh Framework Program 19 (FP7/2007-2013) Universal Influenza Vaccines Secured (UNISEC) consortium under grant agreement no. 602012 and by the EU Horizon 2020 Program under the Marie Skłodowska-Curie grant agreement 713660. DW received a PhD scholarship from the University Medical Center Groningen, Groningen, The Netherlands, and Shantou University Medical College, Shantou, China.

